# Task-specific vision models explain task-specific areas of visual cortex

**DOI:** 10.1101/402735

**Authors:** Kshitij Dwivedi, Gemma Roig

**Affiliations:** Singapore University of Technology and Design, Singapore

## Abstract

Computational models such as deep neural networks (DNN) trained for classification are often used to explain responses of the visual cortex. However, not all the areas of the visual cortex are involved in object/scene classification. For instance, scene selective occipital place area (OPA) plays a role in mapping navigational affordances. Therefore, for explaining responses of such task-specific brain area, we investigate if a model that performs a related task can serve as a better computational model than a model that performs an unrelated task. We found that DNN trained on a task (scene-parsing) related to the function (navigational affordances) of a brain region (OPA) explains its responses better than a DNN trained on a task (scene-classification) which is not explicitly related. In a subsequent analysis, we found that the DNNs that showed high correlation with a particular brain region were trained on a task that was consistent with functions of that brain region reported in previous neuroimaging studies. Our results demonstrate that the task is paramount for selecting a computational model of a brain area. Further, explaining the responses of a brain area by a diverse set of tasks has the potential to shed some light on its functions.

**Author summary:** Areas in the human visual cortex are specialized for specific behaviors either due to supervision and interaction with the world or due to evolution. A standard way to gain insight into the function of these brain region is to design experiments related to a particular behavior, and localize the regions showing significant relative activity corresponding to that behavior. In this work, we investigate if we can figure out the function of a brain area in visual cortex using computational vision models. From our results, we find that explaining responses of a brain region using DNNs trained on a diverse set of possible vision tasks can help us gain insights into its function. The consistency of our results using DNNs with the previous neuroimaging studies suggest that the brain region may be specialized for behavior similar to the tasks for which DNNs showed a high correlation with its responses.

## Introduction

Deep neural networks (DNN) are currently the state of the art models for explaining cortical responses in the visual cortex [1–11]. DNNs trained on a large dataset of images for the object classification task have been shown to explain the human and monkey cortical responses in the inferior temporal cortex (IT) area known for playing a role in object recognition. Further, in Razavi and Kriegsorte [4], it has been revealed that unsupervised models are unable to explain the IT responses as well as the models supervised for the classification task, thus emphasizing that supervision for classification task is crucial for explaining IT responses.

In some recent works [12, 13], they show that the fMRI responses in the scene-selective brain regions like OPA and parahippocampal place area (PPA) are correlated with a DNN trained for the classification task. However, in earlier work from Bonner and Epstein [14], they showed that OPA responses are related to navigational affordances in the scenes. These results raise the question that why a DNN trained for a generic classification task can explain responses for a spatial property like navigational affordances. One possible argument is that the representations of the intermediate layers of a DNN perform generic visual processing. Therefore, these representations are not task-specific and are highly transferrable. An alternative way to explain the above result is that intermediate layers learn representations of the spatial structure of the scene to classify the scene correctly. However, it has not yet been studied in the case of OPA, if a DNN trained for a task related to the function of the brain region in study explains its responses better than a DNN trained on a less related task. We attempt to bridge this gap by taking tasks into account for explaining the brain responses.

In this work, we hypothesize that a DNN trained on a task related to the function of brain region will explain its responses better than a DNN trained on a task which is not explicitly related. Here, we consider that two tasks are different if they generate a different output structure or the predictions are from different domains, for instance, object domain or scene domain. We validate this hypothesis through two different analyses. In the first analysis, we consider a particular case of OPA and explain its responses by a DNN trained on a task (scene-parsing [15]), which we argue is related to navigational affordances. We then compare the results with a DNN trained on generic scene-classification task. In the second analysis, we select DNNs trained on a diverse set of computer vision tasks selected from Taskonomy [16] dataset. We then investigate if the tasks for which DNNs show a high correlation are consistent with functions of scene-selective areas OPA, PPA and Early Visual Cortex (EVC) reported in the previous works [14, 17–23].

The navigational affordances (Fig 1A) as described in Bonner and Epstein [12] is computed by localizing the free space available for navigation in the scene. Thus, a DNN trained on a computer vision task to localize the free space for navigation can serve as a computational model for explaining the navigational affordance related responses in the visual cortex. The scene-parsing (Fig 1B center) task, where the aim is to predict the label of each pixel in the image is, therefore, suitable for our purpose.

**Fig 1.**
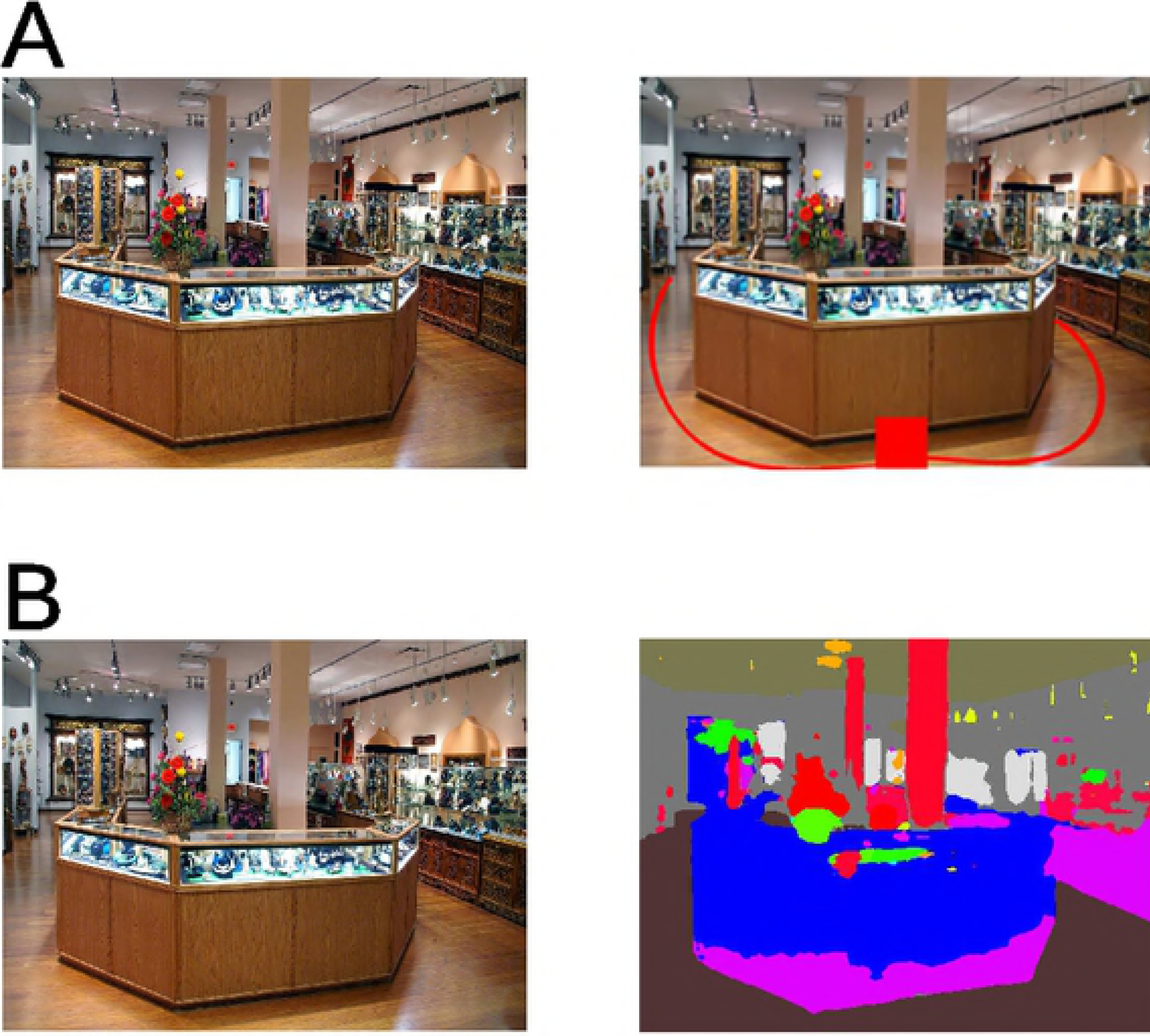
Navigational affordances and scene-parsing task. (A) An example stimulus image (left) presented to the subject for behavior and fMRI experiments in [12]. A path indicated by a rater as instructed in [12] to walk through the scene starting from the bottom center of the image (center). Heat map of possible navigational trajectories produced by combining the data across different raters (right). (reproduced with the permission from Bonner and Epstein [12]). B: Output generated by the scene parsing model (center). Activation of floor unit of the scene-parsing model (right)

The output from the scene-parsing task can label the free space (Fig 1B right) available for navigation and also the obstacles present in the scene. In the scene classification task, the aim is to predict the category of the scene, and the output is a vector containing probabilities of each scene category. Since there is no straight-forward way to localize free space from scene classification output, it is not explicitly related to the navigational affordances. Therefore, we argue that a scene-parsing DNN model will explain the spatial scene property like navigational affordances, and hence, the OPA responses better than a scene classification DNN model.

In the second scenario, we selected DNNs trained on a diverse set of computer-vision tasks from the Taskonomy [16] dataset. The Taskonomy dataset consists of a large set of images with annotations and pretrained DNN models for a diverse set of tasks (Fig 2). We considered all the task DNNs and compared their correlation with OPA, PPA, and EVC. The results of this analysis were consistent with the function of the brain region in consideration.

**Fig 2.**
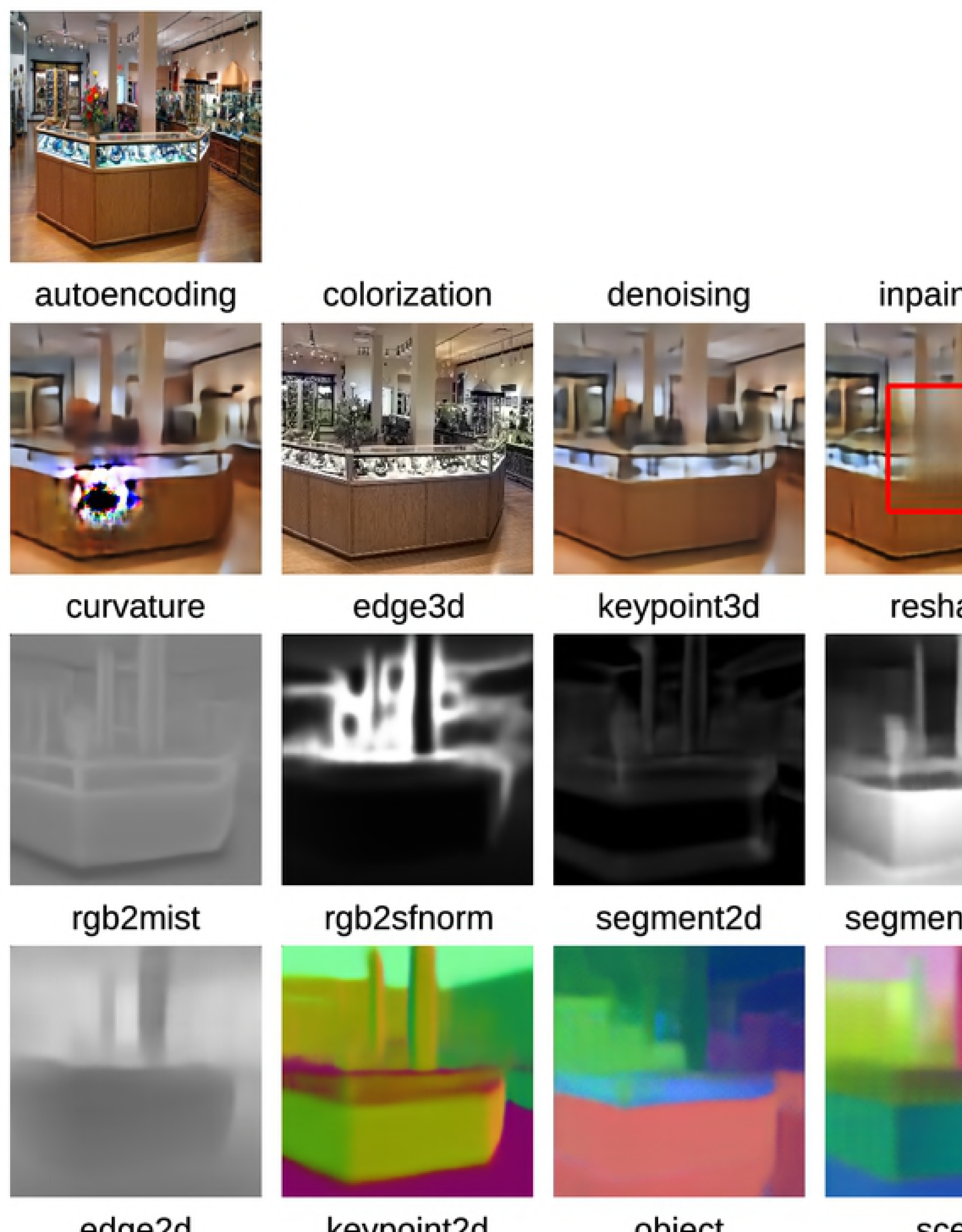
Computer vision tasks from Taskonomy [16] dataset. An example stimulus image (top-left) presented to the subject for behavior and fMRI experiments in [12]. Rest of the images are the output generated by pretrained DNNs optimized for the corresponding tasks selected from the Taskonomy dataset given the stimulus image as the input

Here, we list the key findings of the above analysis:

1. The brain responses of a particular region show high correlation with the DNNs trained on a task related to its function than with the DNNs trained for the classification task.
2. The tasks on which the pretrained DNN activations show high correlation with a particular brain region’s responses are consistent with the functions of the brain region reported in the previous studies.
3. The correlation comparison of a diverse set of task DNN activations with a brain area’s responses provides insight into its previously known/unknown functions.

## Results

In this work, we use Representation similarity analysis (RSA) [24] to compare the correlation of computational and behavioral models with human brain responses. We present the results through two sets of analysis. In the first set, we select a task (scene-parsing) which we argue is similar to the navigational affordances and, hence, the related responses in the OPA. We compute the correlation of brain and behavior Representation Dissimilarity matrices (RDMs) with scene-parsing DNN (VGG_scene-parse_) and compare the results with a scene-classification DNN (VGG_scene-class_). The scene classification task is not as relevant to mapping navigational affordances as the scene parsing task. Therefore, a comparison between these two can provide insights into whether training the DNN in a task related to the function of the brain region in the study is required to explain its responses. In the second set, we select a diverse set of computer vision tasks from the Taskonomy dataset and use DNNs trained for these individual tasks to explain the cortical responses of scene-selective brain regions and early visual cortex. We then compare the correlation with the brain RDMs with the DNNs trained on the above tasks to gain insights into the functions of the brain areas.

### Scene-parsing DNN explains OPA responses related to navigational affordances better than scene-classification DNN

The DNNs are widely used as a potential candidate for computational modeling of areas in visual cortex. Here, we consider two DNNs, one which is optimized on a task (scene-parsing) related to mapping navigational affordances and other on a task (scene classification) not explicitly related to mapping navigational affordances. The DNN models we consider here are VGG_scene-parse_ (Fig 3A) and VGG_scene-class_(Fig 3B) which are similar in architecture except for the last three layers. In VGG_scene-parse_, the last three layers are convolutional to generate spatial mask corresponding to each category, while in VGG_scene-class_, the last three layers are fully connected (FC) to predict the probabilities of possible scene categories. First 13 layers of both the models consist of blocks of convolutional layers with 5 pooling layers in between the blocks.

**Fig 3.**
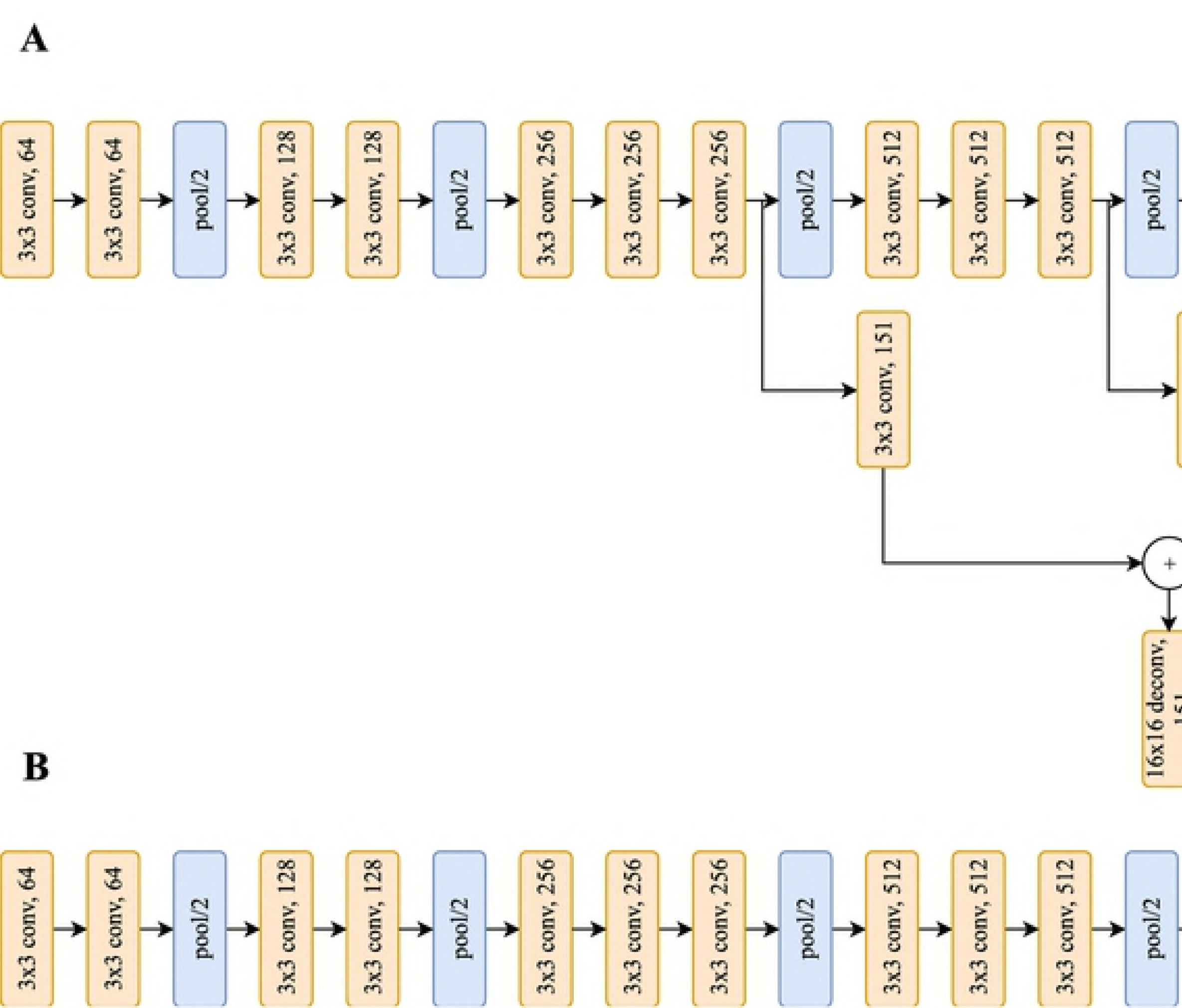
Scene parsing and Scene classification DNNs. (A) DNN model architecture trained on scene-parsing task (VGG_scene-parse_) (B) DNN model architecture trained on scene-classification task VGG_scene-class_. The kxk denotes the kernel dimensions for the convolution and deconvolution layers while the value after the layer type denotes the channel dimensions. Blue layers in the DNNs were selected for comparison

For computing RDMs, we select the DNN activations after the pooling layers and the last three layers of both the DNNs. Fig 4A shows the RDMs of the behavioral model for navigational affordance map (NAM), OPA, and the prefinal layer RDMs of both the DNNs. We argue that deeper layers are more task-related rather than early layers of the DNN. Therefore, we show the RDMs of prefinal layer activations in Fig 4A.

**Fig 4.**
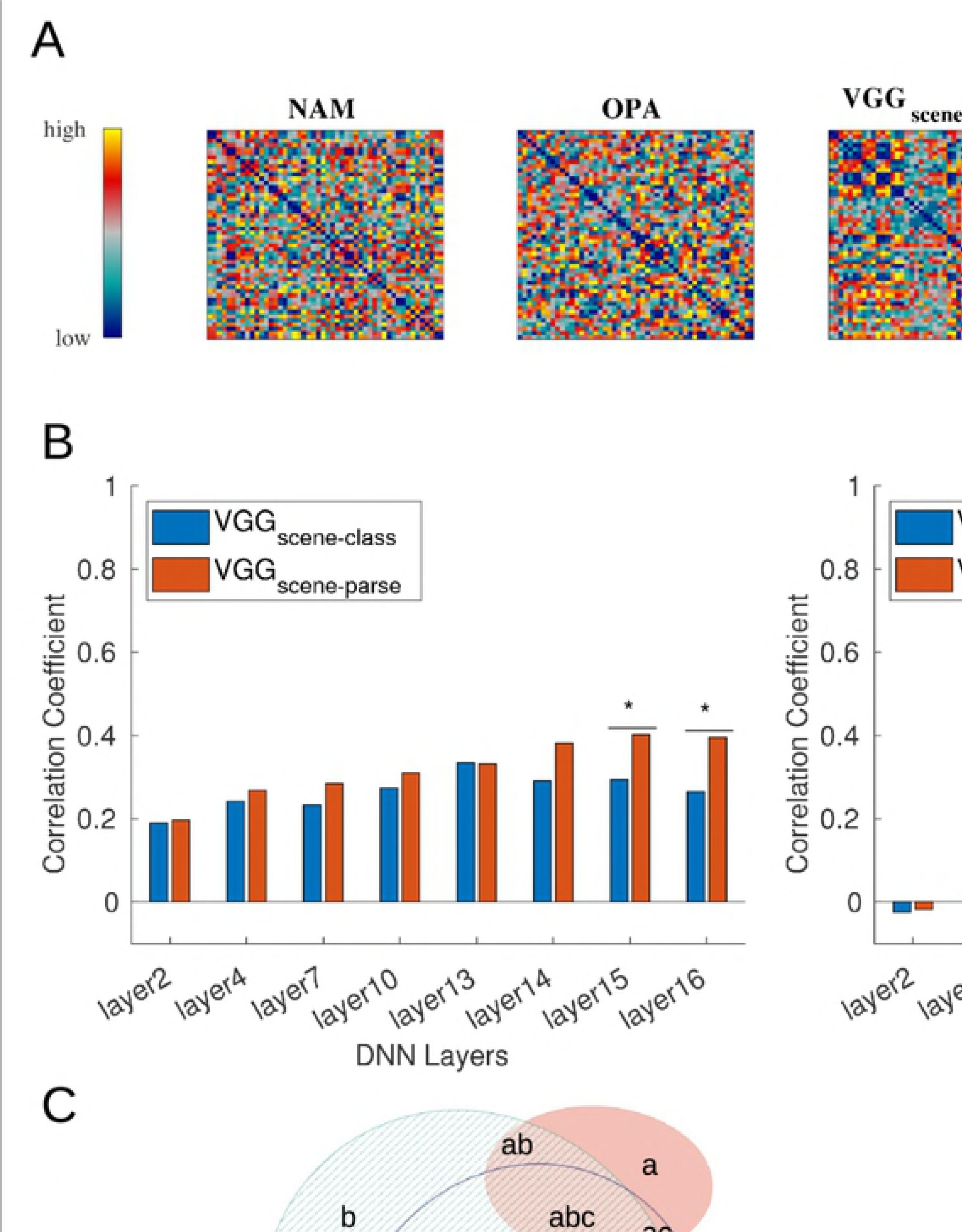
Scene-parsing vs. Scene-classification to explain navigational affordance related responses in the brain. (A) RDMs of navigational affordance, OPA, VGG_scene-class_, and VGG_scene-parse_. (B) (left) Correlation of the OPA responses with VGG_scene-parse_, and VGG_scene-class_, (right) Correlation of the navigational affordance with VGG_scene-parse_, and VGG_scene-class_. (C) Variance partitioning analysis showing the unique and shared variances of behavior and computational models. The asterisk at the top indicates the significance of difference (*p <0.05, **p <0.01, ***p<0.001)

We focus on the correlation with the RDMs of the behavioral model for navigational affordances and OPA responses that are related to navigational affordances. From the results in Fig 4B, we note three key findings:

1. All the layers show a significant correlation (p<0.001) with the brain RDM while only the deeper layers show significant correlation with the behavior RDM
2. Deeper layers for both the DNNs show a higher correlation with brain and behavior RDMs as compared to earlier layers
3. The difference between the correlation values of the deeper layers in both the DNNs with the brain and behavior RDMs is higher and significant (p<0.05 for layer15, layer 16 with OPA, and layer 15 with NAM) in some cases.

The correlation values for all the comparisons are higher and significant for the VGG_scene-parse_ in the deeper layers validating our hypothesis that task-relevant DNNs explain the task-specific regions of the brain better than a generic classification DNN.

### Scene-parsing DNN explains a major portion of the shared variance of the behavior and scene-classification DNN

We combined the RSA with variance partitioning [25] analysis to investigate how uniquely does each model (behavior, VGG_scene-parse_, and VGG_scene-class_) explain the responses of OPA. In variance partitioning approach, using a multiple regression model, we can divide the unique and shared variance contributed by all of its predictors. In this case, OPA RDM was the predictand, and the DNN models and behavior were the predictors. For the DNN models, we selected the RDMs of the layer showing the highest correlation (layer 15 for VGG_scene-parse_, and layer 13 for VGG_scene-class_) with the OPA RDM.

From the results of this analysis (Fig 4C), we note the following points:

1. VGG_scene-class_ shares a major portion (96.62%) of its variance with VGG_scene-parse_.
2. behavior shares more than half of its variance with VGG_scene-parse_ (57.42%) and VGGscene-class (52.35%)
3. VGG_scene-parse_’s unique variance is more than one-forth of the total variance (25.40% of the total variance) explained by the three models

The above results suggest that VGG_scene-parse_ can equally or better account for the navigational affordance related responses in the OPA than VGG_scene-class_, while at the same time uniquely explaining the OPA responses which are neither related to navigational affordances nor scene classification.

### Floor and free space activations explain the behavior but not the brain responses better than the scene-parsing output

The navigational affordance, visually, is related to the free space available for navigation. Therefore, in this analysis, we investigate the case if units corresponding to the free space show a higher correlation with the behavior and brain RDMs than the readout layer of the VGG_scene-parse_. The readout layer of the VGG_scene-parse_ consists of 151 channels with 150 channel each containing an output corresponding to a particular class in the ADE20k [26] dataset and 1 channel corresponding to the background. Therefore, it is straightforward to separate specific category activation from the readout layer. We consider 15 such labels (floor, road, earth, rug, grass, sidewalk, field, sand, stairs, runway, stairway, dirt, land, stage, and step) that represent free space and consider a particular case of floor label as in the dataset by Bonner and Epstein all the stimuli images were from indoor scenes.

We observe from Fig 5A that while floor RDM (*ρ* = 0.2120, p <0.001) showed a higher correlation with the behavior RDM than output RDM (*ρ* = 0.1610, p <0.001), the output RDM (*ρ* = 0.3950, p <0.001) showed a higher correlation with OPA RDM than floor RDM (*ρ* = 0.3054, p <0.001). We perform a similar analysis with PSP_scene-parse_ [27], which is a scene-parsing model that has been shown to achieve higher prediction accuracy than VGG_scene-parse_, to ensure that the results are consistent. The results from Fig 5B show that the trend is consistent among the different models and clears the ambiguity due to the poor performance of the VGG_scene-parse_ model.

**Fig 5.**
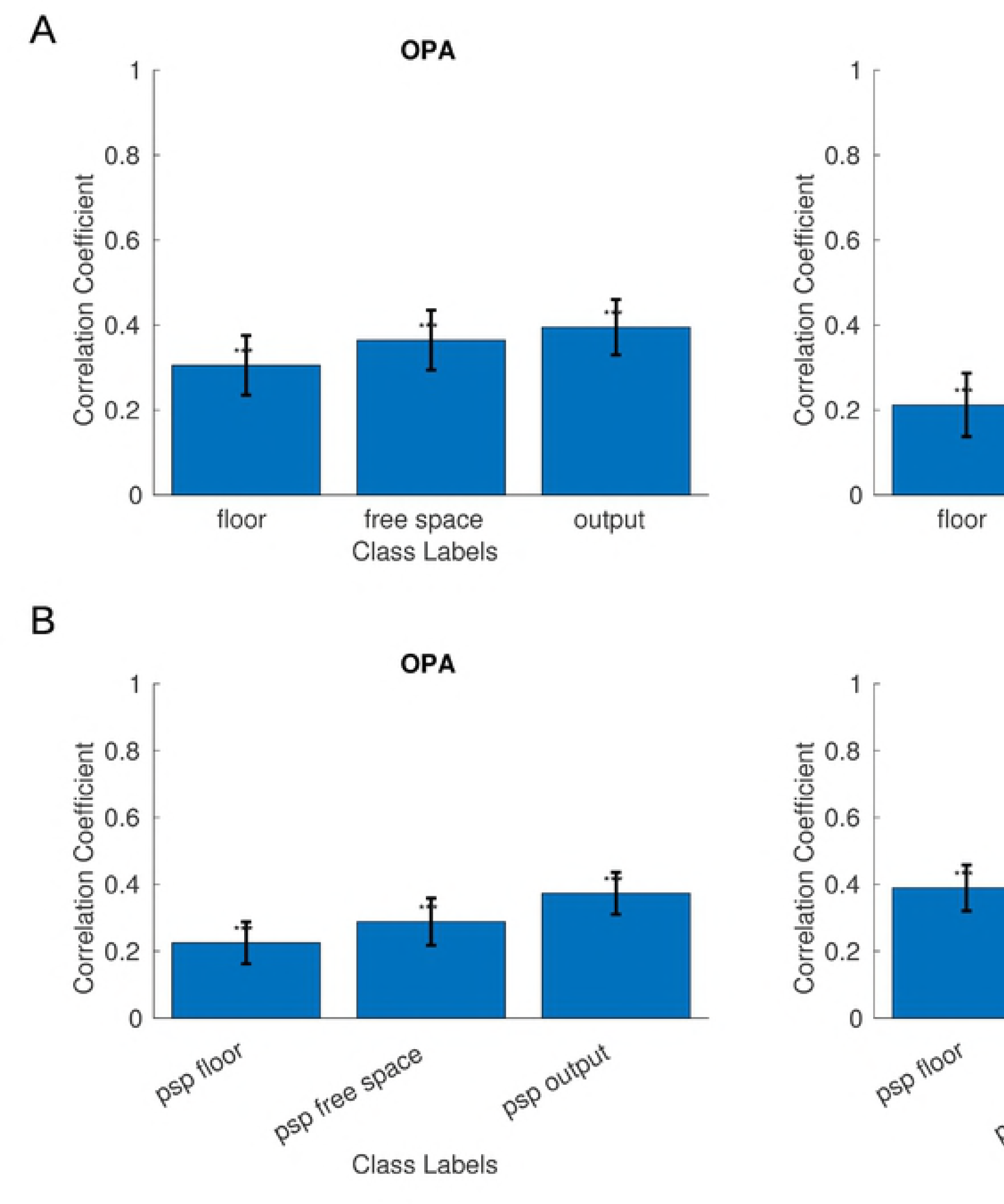
RSA of Scene parsing category-specific activations with OPA and navigational affordances. (A) RSA of floor, freespace and output activations from VGG_scene-parse_ with OPA (left), and behavior (right). (B) RSA of floor, freespace and output activations from PSP_scene-parse_ with OPA (left), and behavior (right). Error bars represent bootstrap *±*1 s.e.m. (*p <0.05, **p <0.01, ***p <0.001)

Further, it is interesting to note that due to more accurate prediction from PSP_scene-parse_ model, the correlation of navigational affordance model with the floor increased to *ρ* = 0.3893 as compared to floor activation from VGG_scene-parse_ (*ρ* =0.2120). The results from this analysis suggest that while just the floor activations can explain navigational affordances, the OPA representation consist of more information about the scene than just navigational affordances.

### Highly correlated categorical DNN units provide insights into the functionality of brain regions

In this analysis, we probe further by computing the correlation of each DNN category unit’s activation with the brain and behavior RDMs. We investigate the top-10 highly correlated categories with the brain and behavior RDMs and observe whether this analysis support the previous works which investigated the functions of OPA and PPA. For this purpose, we use the PSP_scene-parse_ as the predictions generated by PSP_scene-parse_ are more accurate than the VGG_scene-parse_ model.

From Fig 6 (left), we observe that 8 (rug, sidewalk, runway, etc.) out of 10 highly correlated categories with the behavior RDM are indicative of free space with floor RDM showing the highest correlation. Surprisingly, the objects such as vase and clock also showed high correlation with the behavior RDM. A possible explanation for this may be that the vase and clock are generally placed on floor and wall, respectively.

**Fig 6.**
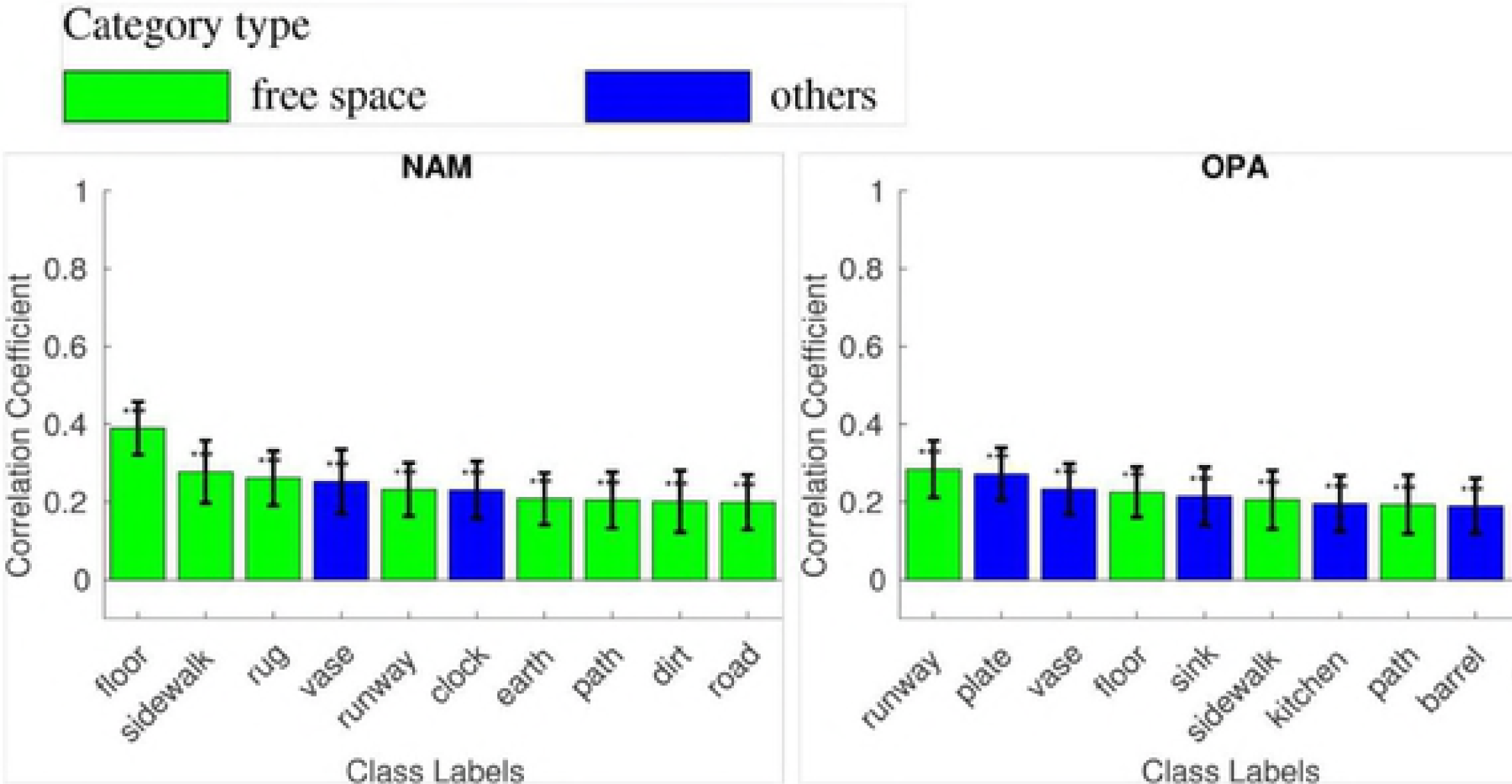
RSA of top-10 correlated categories. RSA of top-10 highly correlated category activations from PSP_scene-parse_ with behavior (left), OPA (center), and PPA (right). Error bars represent bootstrap *±*1 s.e.m. (*p <0.05, **p <0.01, ***p <0.001)

The OPA RDM (Fig 6 center) showed high correlation with only 5 (runway, floor, sidewalk, path, dirt) out of 10 highly correlated categories corresponding to free space. Rest of the labels include object categories: plate, vase, sink, kitchen, and barrel. One possible explanation for these categories being highly correlated is the experimental design [14] in which the OPA responses were recorded. The subjects were asked to classify whether the room displayed is a bathroom or not. The objects such as sink, plate, and vase are highly indicative of the room type, and therefore, OPA responses may be related to the scene classification task. Hence, the high correlation of OPA with these objects is explained by assuming that OPA is involved in the scene classification task. Further, knowing the scene category is also crucial for planning navigation. A related possible explanation is that the objects also suggest the spatial layout of the scene by indicating the presence of obstacles and therefore can be relevant for navigational affordances.

PPA, on the other hand, is hypothesized to represent the spatial layout of the scenes and is insensitive to the navigational affordance as shown in [14]. The results from this analysis (Fig 6 right) are consistent with [14], as the majority of the categories with high correlation are objects that are indicative of scene layout and category and only 2 of the highly correlated categories correspond to free space.

The above analysis demonstrated that categorical units from the scene parsing output are consistent with the functions of the OPA and PPA reported in the previous studies. This result suggests that a categorical analysis has the potential to be used in investigating the functions of brain regions.

### Different task DNNs show different correlation with brain responses

In the above comparison analysis, there were several variables which could play an important role in explaining the brain data. Such variables are the dataset used for learning the DNN models, the DNN architecture, and the tasks for which the DNN was optimized. To disentangle the aforementioned variables, in this analysis, we use the pretrained DNN models which have the same architecture (except the readout layer), and that are optimized on the same set of images from the Taskonomy dataset [16], to perform different tasks.

The provided Taskonomy DNNs architectures consist of an encoder which is same for all the tasks, and a decoder which can vary depending on the task. The encoder architecture for all the tasks is a fully convolutional ResNet-50 [28] with 4 residual blocks and without any pooling layers. The decoder architecture, however, is task-dependent, for example, the decoder of the classification tasks consists of fully-connected layers while the decoder of the tasks in which the output is spatial consist of all convolutional layers. In this analysis, we consider the tasks in which the output is spatial and therefore the decoder architecture is same across all the tasks. In this way, the DNN architecture is the same across all the selected tasks, and only the task is the variable.

We argue that deeper layers of the DNNs decoder may be more task-specific than early layers. To support our argument, we report the mean and variance correlation between the RDMs of several layers from the DNNs and brain RDM in Fig 7. We observe that in earlier layers of the encoder and decoder, the correlation values do not vary significantly across the tasks while the variance increases as we go deeper (Fig 7A left). The mean correlation remains almost constant and decreases on going deeper (Fig 7A center). Yet, the maximum correlation with the brain RDMs consistently increases as we go deeper (Fig 7A right) in the DNN architecture. These results when considered together suggest that for deeper DNN layers, DNNs trained on related tasks show higher correlation while the DNNs trained on unrelated tasks start showing lower correlation values with the brain RDMs. We also observed that the order of tasks correlation values is more consistent in the deeper layers. The above analysis provides evidence that for comparison we should consider the deeper layers of the DNN decoder for all tasks. Thus, in the following experiments, we use the prefinal layer of the decoder for all the tasks to perform the correlation analysis between each of the DNNs RDMs and the brain RDM.

**Fig 7.**
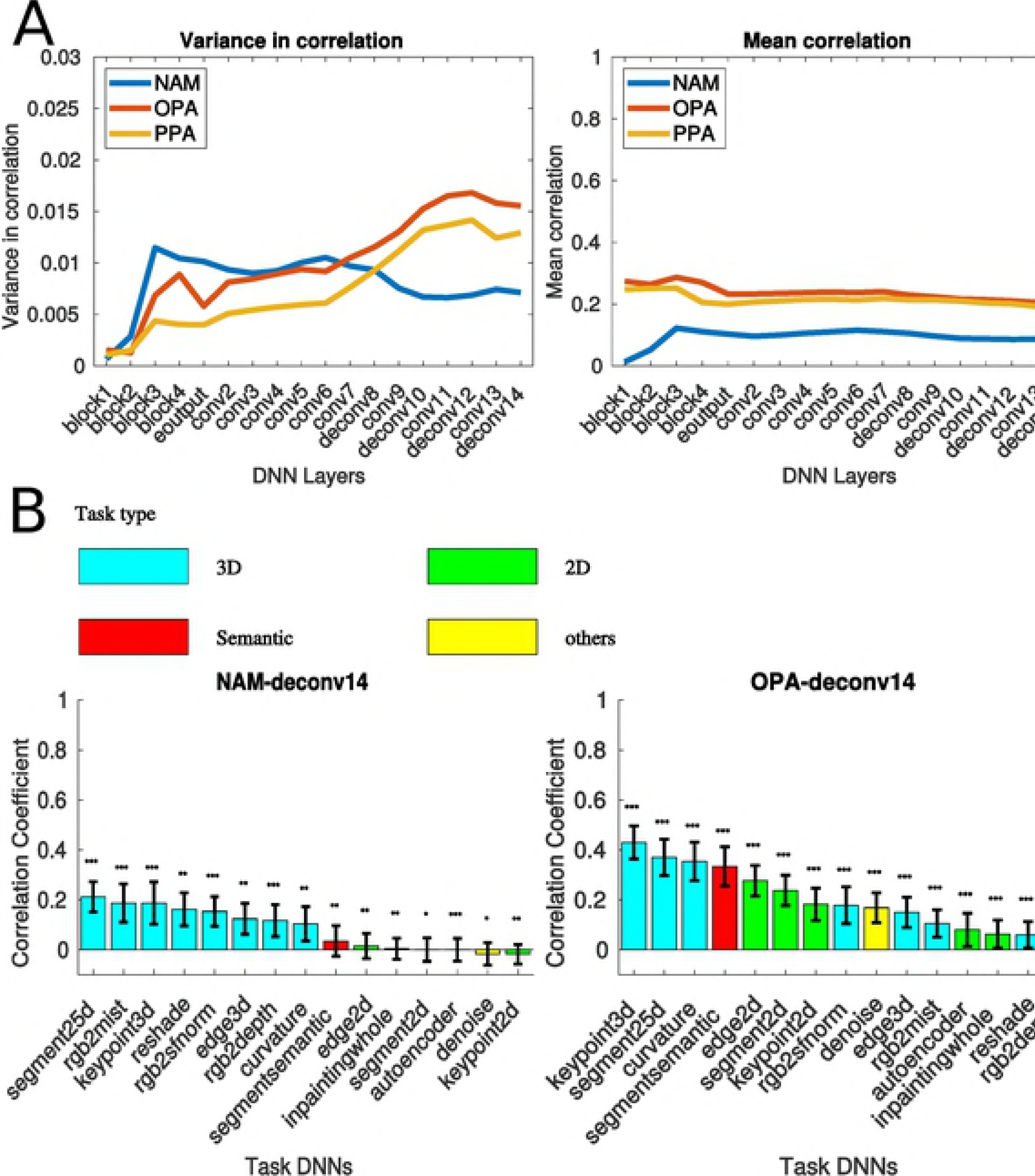
RSA of DNNs (with same architecture) trained on a subset of tasknomy tasks with navigational affordances, OPA, and PPA. (A) Correlation variance (left), mean (center), and maximum (right) of different task DNNs across different layers. The correlation between a task DNN and brain responses was computed for multiple layers located at different depths in the DNN. For the encoder, we selected the output after each residual block and for the decoder we selected the output after each layer. (B) RSA of DNNs (with same architecture) trained on a subset of Taskonomy tasks with behavior (left), OPA (center), and PPA (right). Error bars represent bootstrap *±*1 s.e.m. (*p <0.05, **p <0.01, ***p <0.001)

In this analysis, we focus on the correlation with the RDMs of the behavioral model for navigational affordances, OPA responses that are related to navigational affordances, and PPA responses that are related to spatial layout and scene classification. From Fig 7B, we observe that tasks that are related to 3-dimensional layout of the scene such as 3-d keypoints (*ρ* = 0.4299 for OPA, and *ρ* = 0.4253 for PPA), 2.5d segmentation (*ρ* = 0.3702 for OPA, and *ρ* = 0.3609 for PPA) and curvature (*ρ* = 0.3543 for OPA, and *ρ*= 0.2816 for PPA) show the highest correlation with both the OPA and PPA RDMs. Further, the semantic segmentation task (*ρ* = 0.3341 for OPA, and *ρ* = 0.2652 for PPA) that requires categorical and spatial layout information also shows a high correlation with both these brain regions. On the other hand, tasks such as autoencoding (*ρ* = 0.0804 for OPA, and *ρ* = 0.0715 for PPA), and inpainting (*ρ* = 0.0630 for OPA, and *ρ* = 0.0527 for PPA) that are neither related to 3-dimensional representation or scene-semantics show low and insignificant correlations with the brain areas. However, on comparing with behavior RDM, we observed that 2.5-D segmentation (*ρ* = 0.2122) although showed the highest correlation but semantic segmentation task showed low correlation (*ρ* = 0.0352). We argue that the low correlation of semantic segmentation task DNN with behavior is because the categories in this task do not contain floor or free space as compared to scene-parsing task.

The results from the above analysis suggest that for explaining the brain responses of task-specific brain regions using the DNNs, the DNN should be optimized on a task related to the function of that brain region. The above analysis also reveals that DNNs of same architecture that are trained with the same dataset show a difference in correlation due to the task.

### Task optimized DNNs provide insights into the functionality of brain regions

Above, we demonstrated through a series of analysis that a DNN optimized to perform a task related to a particular brain region function, better explains the responses of such brain region. In this analysis, we take a different route and ask the question if we can gain an insight about the functions of the brain region by comparing the correlation of different task DNNs with the brain RDMs.

In the current analysis, we focus on the correlation with the RDMs of OPA, PPA, and EVC. We consider all the single image tasks from the Taskonomy dataset and use the prefinal layer RDM as the representative DNN RDM for that particular task. Then, we perform the comparison of correlation between the brain RDMs and the prefinal layer RDMs of the DNNs trained on selected tasks.

The results (Fig 8A) of OPA and PPA are similar to the previous analysis. Now, we also observe a high correlation for both areas and the scene and object classification tasks, which were not reported in the previous analysis. These results suggest that while PPA and OPA representations are spatial to perform tasks requiring spatial layout information, the representations in these areas are also semantic to perform classification tasks. We observe that EVC is highly correlated to edge2d (*ρ* = 0.6972, p = 0.0002), object classification (*ρ* = 0.6289, p = 0.0002) while insignificantly related to tasks such as vanishing point (*ρ* = 0.0516, p = 0.0546) and rgb2mist (*ρ* = 0.0567, p = 0.0414).

**Fig 8.**
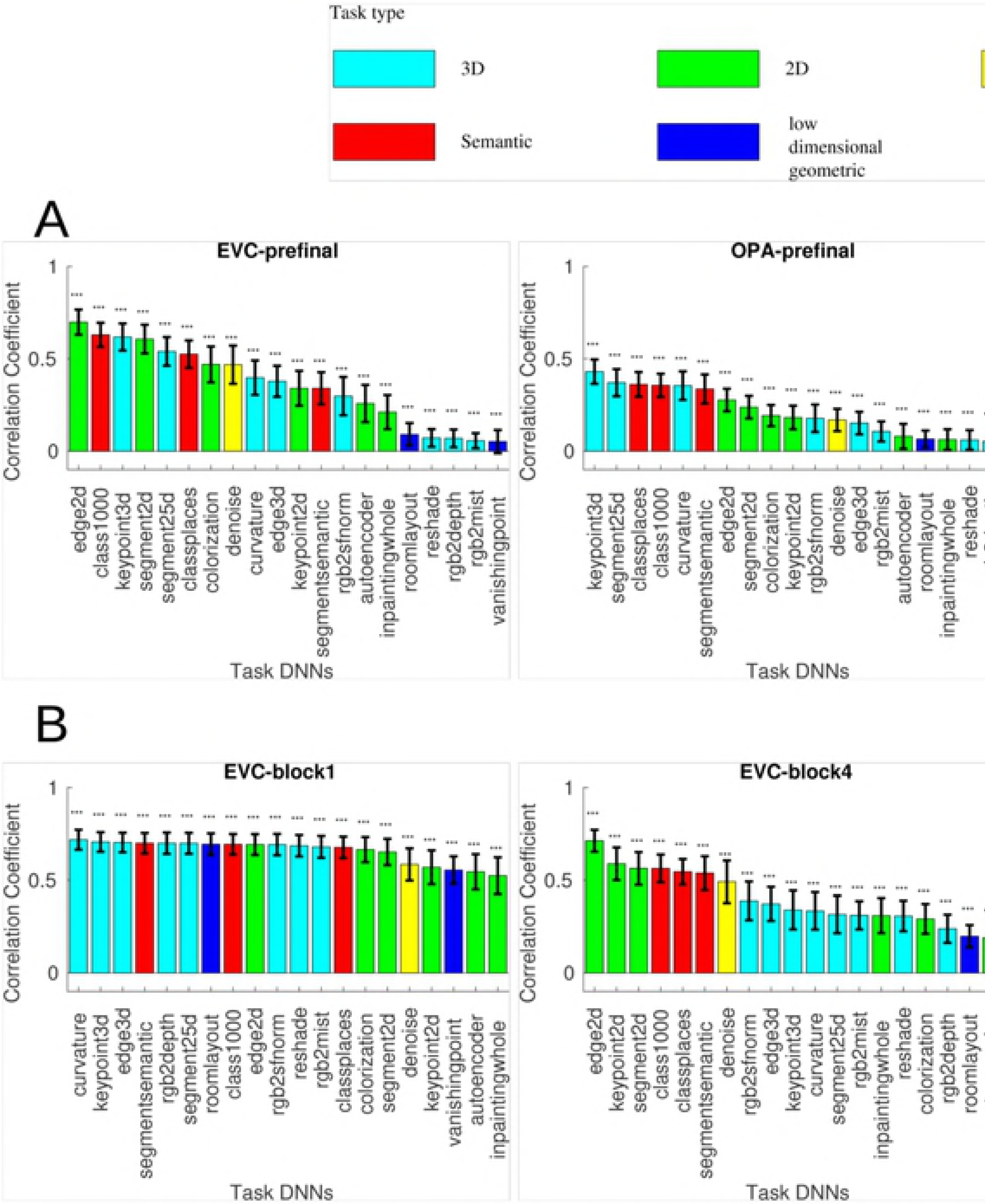
RSA of DNNs trained on tasknomy dataset with EVC, OPA, and PPA. (A) RSA of DNNs trained on Taskonomy tasks with EVC (left), OPA (center), and PPA (right). (B) RSA of three layers of DNNs at early stage (left), middle stage (center), and final stage (right) trained on a subset of Taskonomy tasks with EVC. Error bars represent bootstrap *±*1 s.e.m. (*p <0.05, **p <0.01, ***p <0.001)

The EVC and the initial layers of the DNN are known to have a general visual representation that is not task-specific while the deeper layers and regions of higher visual cortex are known to have more task-specific representations. To investigate this, we probe further into EVC by observing the correlation of EVC with different task DNNs (Fig 8B) at block 1, block 4 layers of the encoder and prefinal layer of the decoder. We observe that in block 1 all the tasks have very high and significant correlation (*ρ>*0.5, p<0.001) with the EVC RDM. On the other hand, the correlation for some tasks starts dropping as we move from block 1 to block 4 and prefinal layer. The results support previous work [7] showing that EVC representation is more similar to early layers of DNN. Further, the tasks which show very high correlation with the EVC in deeper layers are mostly related to low-level visual cues (edge2d, keypoint 3d, segment2d, etc.) or the classification (object and scene classification). The high correlation with the classification DNNs may be due to the emergence of object detectors in the early visual cortex similar to as shown to emerge in DNNs [29].

Thus, the above analysis shows that performing RSA of a brain region with a diverse set of tasks has the potential to shed some light on the functionality of that particular brain region in the visual cortex.

### Similar task DNNs share more variance than dissimilar task DNNs

We probe further whether the tasks that are similar according to the Taskonomy transfer matrix share more variance than the tasks that are less similar. This analysis is performed to investigate whether two dissimilar tasks can be used to uniquely explain the responses of brain areas corresponding to different behaviors.

We use the variance partitioning [25] approach to calculate the unique and shared variance of different models. The brain RDMs (OPA, PPA, and EVC) are the predictand, and three task DNNs (pairs of similar and dissimilar tasks) are the predictors. We used the RDMs of the prefinal layer for all the DNNs tested.

Following Fig 8A, the tasks selected in the similar pair for OPA were Keypoint3d and curvature, for PPA were Keypoint3d and curvature, and for EVC were edge2d and segment2d. The tasks selected in the dissimilar pair for OPA were Keypoint3d and scene-classification, for PPA were Keypoint3d and scene-classification, and for EVC were edge2d and object classification.

The results from partitioning analysis (Fig 9A) of all three brain RDMs show that unique variance of dissimilar task (19.93% vs. 4.13% for OPA, 18.73% vs. 1.35% for PPA, 6.53% vs. 2.10% for EVC) is higher than the similar task. This analysis suggests that two DNNs optimized for dissimilar tasks may be used to explain the brain responses uniquely related to each task.

**Fig 9.**
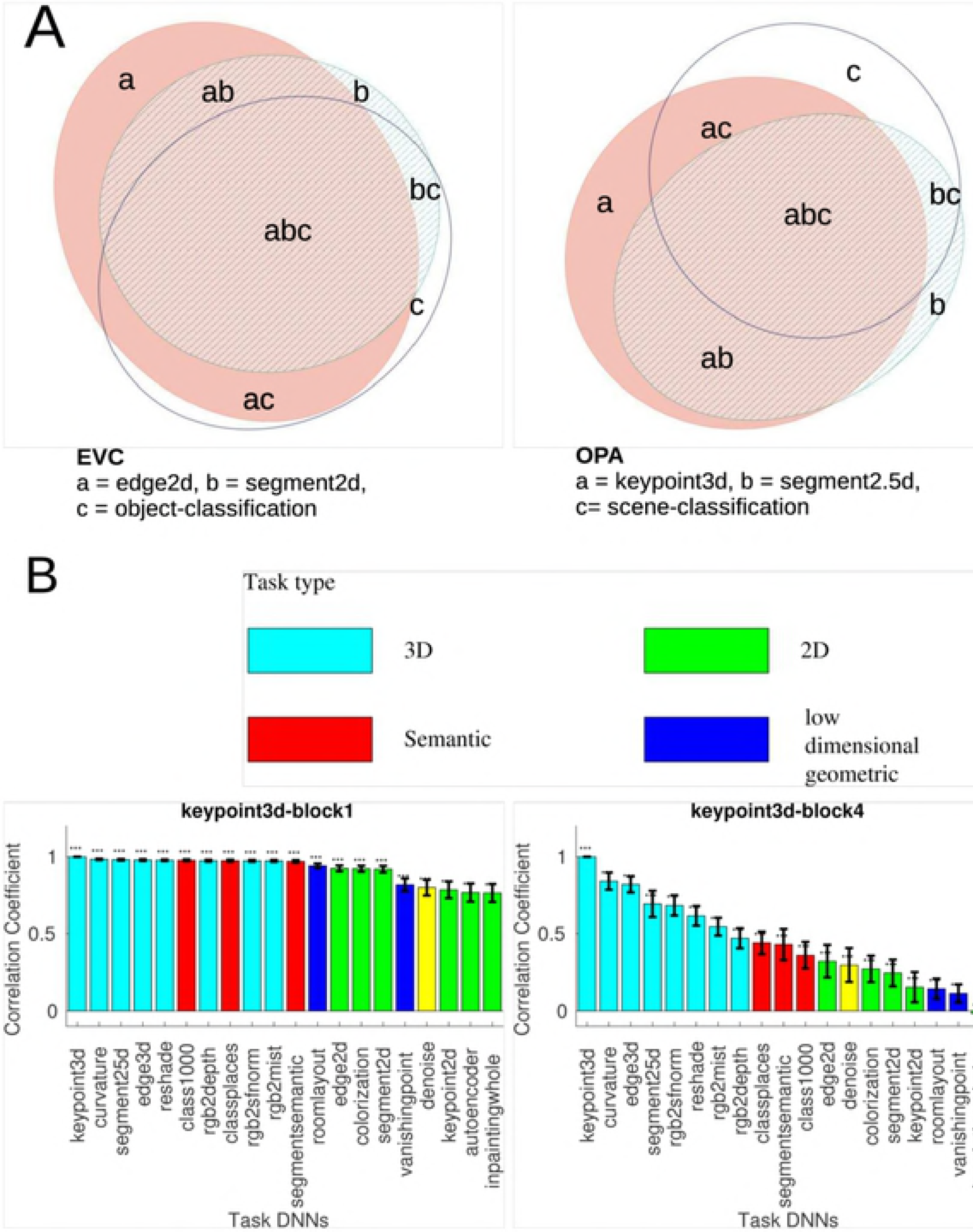
Task similarity using variance partitioning analysis and RSA. (A) Variance partitioning analysis of EVC (left), OPA (center), and PPA (right) with three task DNNs. Tasks a and b are similar while tasks a and c are dissimilar for all the three cases. (B) RSA of keypoint3D DNN with other task DNNs at the early stage (block 1, left), center stage (block4, center), and final stage (prefinal layer, right). Error bars represent bootstrap *±*1 s.e.m. (*p <0.05, **p <0.01, ***p <0.001)

### Correlation between different task DNNs

In this analysis, we investigate how correlated are the DNN representations across different tasks. For this purpose, we choose a single task (3D keypoints) DNN and compute the correlation with other task DNNs across different layers in the DNN architecture. First observation (Fig 9B left) to note is that earlier layers of all the tasks are highly correlated with each other. Secondly, in the deeper layers (Fig 9B center, right) the task similar according to Taskonomy transfer matrix show high correlation whereas the dissimilar tasks show a lower correlation with the task in consideration.

## Discussion

In this work, we demonstrated the importance of task selection for using pretrained DNNs as a computational model for task-specific regions of visual cortex. We list the key findings from our analyses below.

- A DNN trained on a task (scene-parsing) related to the function (navigational affordance) of the brain region (OPA) shows a higher correlation with its responses than a DNN trained on a task (scene-classification) not explicitly related.
- Category-specific activations generated from the scene-parsing DNN provide insights into functions of the scene-selective cortex.
- Training DNNs with same architecture on the same dataset but for different tasks results in different correlation with the brain responses.
- DNNs that show a high correlation with the brain responses are trained on tasks related to the functions of the brain areas reported in the previous studies.

In the following paragraphs, we discuss the strength and limitations of the key analysis and findings of this work.

### Finding a DNN trained on a task related to the function of the brain region

In our first analysis, we selected a scene-parsing task and showed how it could be related to navigational affordances in the scene. However, it is not always possible to find a computer vision task that could be explicitly related to the function of the brain area. Further, even if we find a related task, there might be a possibility that the annotations for such a task are not readily available. Hence, finding a DNN pretrained on this particular task might not be possible. Likely, due to these reasons most of the previous studies adhered to a DNN pretrained on classification.

With the advance in the computer vision field, new datasets with annotations are continuously made available to the public. The Taskonomy dataset used in this work is a large-scale image dataset with annotations available for most of the commonly studied computer vision task on images. For more complex tasks, such as navigation planning, and visual question answering there are new datasets [30–34] available with virtual 3D environments where an agent can navigate and interact with the environment. The DNN models trained on these datasets can shed some light on the functions of brain areas related to visual memory, navigation, and interaction with the environment. Thus, as new datasets for a wide variety of tasks, and hence, pretrained DNNs on these datasets become available, the comparison of these DNNs with the brain responses will shed new light into the functions of different brain regions. Through a specific example of OPA, navigational affordances, and scene-parsing task, we believe our work has paved a way towards future studies using task-optimized DNNs as potential computational models for task-specific brain regions.

### RSA with categorical units: a potential method to investigate functions of a semantic brain area

We showed that the responses of categorical units which are generated as the output of the scene-parsing task could be used to gain insights into the functions of OPA and PPA. For OPA, while half of the top correlated category units corresponded to free space for navigation, other half corresponded to objects indicative of scene category.

The result was consistent with earlier neuroimaging works investigating the function of OPA [14, 35, 36]. While in Bonner and Epstein [14], it is shown that OPA is involved in navigational affordance of scenes, in Dilks et al. [36] it has been shown that OPA might also be playing a role in scene classification. Similarly, for PPA, the top correlated classes mostly contained objects indicative of scene category and layout and only 2 out of the top 10 correlated classes corresponded to free space. These results further provide the evidence that PPA responses are insensitive to navigational affordance which is also consistent with the findings related to PPA in Bonner and Epstein [14]. Thus, the above results suggest that RSA with categorical activations can be a potential method to investigate functions of a brain area. Further, it is important to note that we were unable to distinguish the functions of OPA and PPA through the analysis involving a diverse set of tasks since both the OPA and PPA RDMs showed high correlation with the same set of tasks. In such cases, where the functional difference is due to semantical categories and not the spatial tasks, the categorical activations can be used to distinguish the functions of these brain regions.

However, there are few potential shortcomings with this approach. First, the number and type of categories are limited by the dataset used for training the DNN. Therefore, in a new set of stimuli which contains categories that were not present in the dataset used for training of DNN, the top correlated classes might not provide any useful insights. Also, it is not always the case that the brain areas are categorical and therefore this approach may not provide any meaningful insight into the functionality of those brain areas.

### Difference in correlation: Is it because of task?

In the first analysis, we found that scene-parsing DNN showed a higher correlation with OPA and navigational affordances than the scene-classification DNN. Yet, it is important to note that there were three differences in the DNNs used for comparison. First was the architecture difference, while the last 3 layers of scene-parsing DNN were convolution, the last 3 layers of scene-classification DNN were fully connected. Second, the dataset used for training both the DNNs were different, ADE20k [15] for scene-parsing DNN and Places-365 [37] for scene-classification DNN. Thirdly, the task on which the models were trained were different. Therefore, the difference in the correlation of 2 DNNs with OPA and navigational affordances could be attributed to any of these factors. To clear this ambiguity, we selected DNNs with same architecture trained on the same set of images but for different tasks. We then showed that the DNNs trained on tasks related with the function of brain area were highly correlated while the DNNs trained on unrelated tasks showed low or insignificant correlation with the brain area. From this analysis, we found that training on different tasks leads to a difference in correlation of the DNN activations with the responses of the brain regions. The correlation of the DNN with the brain region depends on how similar the task is with the function of the brain region.

### DNNs trained on a diverse set of tasks: a potential method to assess unknown functions of a brain area

We compared the correlation of DNNs trained on a diverse set of tasks with different brain areas. The above comparison was performed to investigate whether the highly correlated tasks are related to and are consistent with the previously reported functions of the brain areas. The top-3 task DNNs (3D Keypoints, curvature, and 25d segmentation) showing the highest correlation with the scene-selective visual areas (OPA and PPA) were related to the 3-D structure of the scenes. In an electrophysiological study [21], they demonstrated the importance of structure defining contours through the electrophysiological investigations of scene-selective visual cortex in the macaque brain. In a related neuroimaging work [22], they showed that scene category could be decoded from the PPA even if the stimuli images are just the line drawings of the corresponding scene. In Choo and Walther [23], they show that intact contour junctions are crucial for scene category representation in PPA. Thus, the high correlation of OPA and PPA responses with the DNNs trained to predict 3-D keypoints and curvature demonstrate that our results are consistent with the previous studies investigating the representation of scene-selective visual cortex.

Further, the semantic tasks such as scene/object classification and semantic segmentation also showed high correlation with the scene-selective visual cortex. This is consistent with the results of Bonner and Epstein [14] and Epstein et al. [17] where they report that representation of OPA and PPA is also semantic. Thus, the results from the task comparison analysis are consistent with the previous studies of the scene-selective visual cortex and provide the evidence that the representation of scene selective visual cortex is both semantic and visual. Further analysis with the early visual cortex showed that the tasks which require low-level vision cues such as 2d edges, 2d segmentation are highly correlated with the EVC responses. These results taken together suggest a strong potential for using a diverse set of tasks for gaining insights into the function of different brain regions.

One counter-argument to our approach might be that humans are never supervised the same way as these DNNs. The DNNs were supervised using the task-specific annotations. However, no such annotations are available to humans, and they learn to perform these tasks intuitively. However, one should also note that humans learn through interaction with the environment, by moving around, and learning from others. Therefore, these intermediate vision tasks may have been learned through supervision of much complex goal. Learning complex tasks is still a challenging area in the artificial intelligence and a single model is not able to perform all the tasks a human can perform. Therefore, in this work, we focused only on scene-selective regions in visual cortex and tried to explain its responses by DNNs trained on different tasks. Further, in this work, we are only interested in the correlation of the end state of the representations and not how either the DNNs or the human learned these representations.

## Conclusion

In this study, we presented the evidence supporting our hypothesis of using task-specific DNN models to explain responses of task-specific brain regions. We first validated this hypothesis by considering the particular case of OPA which has been reported to be associated with navigational affordances. We showed that a scene-parsing DNN that is related to the navigational affordances shows a higher correlation with OPA responses than a DNN trained on a less related task (scene-classification). We further validated this hypothesis by comparing the correlation of the responses of scene-selective visual areas with a large and diverse set of task DNNs. Although in this work, we only considered scene-selective visual areas we believe that the similar results can be obtained for other higher cognitive brain areas such as hippocampus and prefrontal cortex. One other limitation of this work is that we only considered the tasks that apply to single static images. In future studies, we aim at considering performing a similar analysis with more complex functions and with models trained on complex tasks in virtual 3-D environments. We believe this study has paved a way towards using task-optimized DNNs as potential computational models for task-specific brain regions.

## Materials and methods

In the first section, we describe Representation similarity analysis (RSA) [24] which is a standard method to compare the correlation of computational and behavioral models with human brain responses. In the second section, we describe the variance partitioning analysis which was used to find the unique and shared variance of the computational models used to predict brain responses. In the third section, we briefly describe the dataset we used in this work and then in the last section we provide the details of the DNN models used for analysis.

### Representation similarity analysis (RSA)

RSA is used to compare the information encoded in brain responses with a computational or behavioral model by computing the correlation of the corresponding Representation Dissimilarity matrices (RDMs). In the case of comparison with DNNs, we compute the correlation of RDMs of the brain responses with the RDM of layer activations of the DNNs.

#### Representation Dissimilarity Matrix (RDM)

The RDM for a dataset is constructed by computing dissimilarities of all possible pairs of stimulus images. For fMRI data, the RDMs are computed by comparing the fMRI responses while for DNNs the RDMs are computed by comparing the layer activations for each image pair in the dataset. The dissimilarity metric used in this work is 1*-ρ* where *ρ* is the Pearson’s correlation coefficient. Although in the previous work [12], where a scene classification DNN was compared with the navigational affordance the dissimilarity metric used was the Euclidean distance, we observed that with 1*-ρ* as the dissimilarity metric, the correlation was higher. Hence, in this work for all the analysis 1*-ρ* is used as the dissimilarity metric to compute RDMs of layer activations. We did not perform PCA on layer activations as done in [12] since the spatial information in the case of convolutional layer outputs is lost. The spatial information is lost because for performing PCA as done in [12]; first, the convolutional layer output is flattened and then principal components are selected due to which information from some spatial location is never considered for the analysis. For the first set of analysis with scene-parsing and scene-classification DNN, we consider OPA and PPA RDMs for comparison as these areas have been hypothesized to represent scene affordances [14] and scene layout [17] respectively. We also compare the DNN RDMs with a behavior Navigational Affordance Map (NAM) [12] that represents navigational affordances in a scene. For the second set of analysis with Taskonomy DNNs, we consider OPA, PPA, and EVC RDMs to compare with DNN RDMs.

#### Statistical analysis

We use RSA toolbox [38] to compute RDM correlations and corresponding p-values and standard deviation, using bootstrap similar to [12]. For determining which RDM better explains the behavioral or neural RDMs, we perform a two-sided statistical comparison. The p-values are estimated as the proportion of bootstrap samples further in the tails than 0. The number of bootstrap iterations for all the analysis was set to 5000.

### Variance partitioning analysis

Variance partitioning method is used to determine the unique and shared contribution of individual models when considered in conjunction with the other models. We describe the analysis by considering the case of OPA predicted by behavior model related to navigational affordance, scene-parsing DNN, and scene-classification DNN. First, the off-diagonal elements of the OPA RDM is assigned as the dependent variable (predictand). Then, the off-diagonal elements of behavior RDM and layer RDMs representing scene-parsing and scene-classification tasks are selected as the independent variable. Then, we perform seven multiple regression analysis: one with all three independent variables as predictors, three with three different possible combinations of two independent variables as predictors, and three with individual independent variables as the predictors. Then, by comparing the explained variance (*r*^2^) of a model used alone with the explained variance when it was used with other models, the amount of unique and shared variance between different predictors can be inferred. For the other variance partitioning analysis in this work, the predictors and predictands were modified accordingly, and the steps of analysis were the same. The area proportional venn diagrams for the variance partitioning analysis were generated using EulerAPE software [39].

### Navigational Affordance Dataset and Model

The stimuli images used for analysis consisted of 50 images of indoor environments. The subject’s fMRI responses were obtained while they performed a category-recognition task (bathroom or not). In this work, we directly use the precomputed subject averaged RDMs of the navigational affordance map (NAM), PPA and OPA provided by Bonner and Epstein [12].

To obtain NAM, first, an independent group of subjects was asked to indicate the paths in each image starting from the bottom using a computer mouse. The probabilistic maps of paths for each image were created followed by histogram construction of navigational probability in one-degree angular bins radiating from the bottom center of the image. This histogram represents a probabilistic map of potential navigation routes from the viewer’s perspective. For further details of the navigational affordance map or dataset, please refer to [12, 14].

### Deep Neural Network Models to explain brain responses

In this section, we describe the architecture of the DNN models used in the analysis.

#### Scene-classification model

We choose VGG16 [40] trained on Places [37] (a scene classification dataset) dataset as the scene classification model (VGG_scene-class_) (pretrained model downloaded from https://github.com/CSAILVision/places365). The reason behind the different choice of scene classification model from the one used in Bonner and Epstein [12] was that we were unable to find a pretrained scene-parsing model with similar architecture as Alexnet [41]. VGG16 model (Fig 3A) contains 13 convolutional layers with 5 pooling layer after a convolutional block of either 2 or 3 convolutional layers and 3 fully connected (FC) layers after the last pooling layer.

#### Scene-parsing models

We use fully convolutional modification of VGG16 [42] trained on ADE20k [26], [15] (a scene-parsing dataset) as the scene-parsing model (VGG_scene-parse_). In VGG_scene-parse_ (pretrained model downloaded from https://github.com/hellochick/semantic-segmentation-tensorflow) (Fig 3B), the FC layers are replaced by convolutional layers to predict pixel-wise spatial mask. The model has additional deconvolutional layers to upsample the spatial mask obtained from the intermediate layers. We use pyramid scene parsing network (PSP_scene-parse_) for performing analysis of category specific outputs as PSP_scene-parse_ (pretrained model downloaded from https://github.com/hellochick/semantic-segmentation-tensorflow) outperforms VGG_scene-parse_ on scene-parsing task and hence the categorical outputs are more accurate and suitable for that particular analysis. The PSP_scene-parse_ model introduces a pyramid pooling module that fuses features of four different scales to obtain superior performance over VGG_scene-parse_.

#### Taskonomy models

Taskonomy dataset is a large-scale image dataset containing 4 million images with annotations and pretrained DNN models available for 26 vision related tasks. The tasks included in this dataset cover most common computer vision tasks related to 2D, 3D, and semantics. The tasks involved range from low-level visual tasks like edge detection to more abstract semantic tasks like scene/object classification. The architecture of DNNs (pretrained models downloaded from https://github.com/StanfordVL/taskonomy/tree/master/taskbank) trained on different tasks from Taskonomy dataset share a common encoder architecture. The encoder is a fully convolutional ResNet-50 [28] without any pooling layers. The encoder architecture consists of 4 residual blocks each containing multiple convolutional layers. The decoder architecture, however, varies according to the output structure of each task. For the tasks where the output is spatial (like edge detection, semantic segmentation), the decoder is a 15-layer fully convolutional network. For the tasks where the output is lower dimensional like scene/object classification, the decoder contains 2-3 FC layers.

## Acknowledgments

We thank Michael Bonner for providing the permission to use the images from his paper, and insightful discussions. We thank Astha Gupta and Debidatta Dwibedi for helpful comments on the manuscript.

